# Parameter estimation in the Montijano-Bergues-Bory-Gompertz stochastic model for unperturbed tumor growth

**DOI:** 10.1101/2024.09.09.611959

**Authors:** Beatriz Bonilla-Capilla, Luis Enrique Bergues Cabrales

## Abstract

Different sources of noises endogenous and exogenous to the cancer are involved in its stochastic growth. The aim of this study is to propose the stochastic version of Montijano-Bergues-Bory-Gompertz equation for the unperturbed tumor growth kinetics. The maximum likelihood estimators for the intrinsic tumor growth rate and the growth decelerating factor, and their respective discrete time approximations were analytically calculated. Different simulations of the deterministic and stochastic of this equation were made for different values of their respective parameters. Limit conditions for the average diffusion coefficient and the growth decelerating factor were established. The tumor volume at the infinite was calculated for several values of parameters of the stochastic Montijano-Bergues-Bory-Gompertz equation. Furthermore, descriptive statistic for the maximum likelihood estimators of the intrinsic tumor growth rate was computed for several parameters of this equation. The results evidenced that solid tumors there are for values of the average diffusion coefficient and the growth decelerating factor less than their respective limit values. The transition between avascular and vascular phases of the unperturbed tumor growth kinetics was revealed in the plot of the discrete time approximation for the maximum likelihood estimator of the growth decelerating factor versus the discrete time approximation for the maximum likelihood estimator of the intrinsic tumor growth rate. These results were connected with different findings in the literature. In conclusion, the stochastic Montijano-Bergues-Bory-Gompertz equation may be applied in the experiment to describe the unperturbed tumor growth kinetics, as previously demonstrated for its deterministic version, in order to estimate the parameters of this equation and their connection with processes involved in the growth, progression and metastasis of unperturbed solid tumors.

**Author summary:** In order to comprehend the unperturbed tumor growth, we investigate a new mathematical model called the stochastic Montijano-Bergues-Bory-Gompertz equation. This study is made based on the ideas of Ferrante et al. and the deterministic version of the Montijano-Bergues-Bory-Gompertz equation. By applying this stochastic equation, we aim to provide valuable insights into how tumors grow and spread throughout the body. We focus on estimating key parameters that are essential for understanding the dynamic processes involved in the unperturbed tumor behavior. Our findings may help researchers to understand the stochastic nature of the unperturbed tumor growth; know the existence of transitions in the unperturbed tumor growth kinetics, probably between avascular and vascular phases; and reveal the values of the model parameters for which the solid tumor is functional, non-functional or does not exist. These aspects may be relevant to propose an individualized anticancer therapy aimed at minimizing the different noise sources that occur during the unperturbed tumor growth. Overall, this study contributes to our ongoing efforts to improve cancer treatment strategies and enhance patient outcomes by fostering a better understanding of tumor biology.

## Introduction

Knowledge the unperturbed tumor growth kinetics (TGK) allows us to reveal and understand intrinsic processes in the tumor. This is of vital importance to propose individual anticancer therapies aimed at achieving complete remission of cancer or turning it into a controlled chronic disease.

Different deterministic mathematical models have been used to describe unperturbed TGK, such as: exponential, conventional Gompertz (CG), Logistics and Bertalanffy-Richards equations, being the deterministic CG equation better describes experimental data [1, 2]. Furthermore, the deterministic equations of Kolmogorov-Johnson-Mehl-Avrami (KJMA) and modified Kolmogorov-Johnson-Mehl-Avrami (mKJMA) [3], and Montijano-Bergues-Bory-Gompertz (MBBG) [4] have been suggested to fit the unperturbed TGK.

The deterministic KJMA, mKJMA and MBBG equations do not differ significantly from the deterministic CG equation when their fits are compared [3, 4]. In a previous study, the good correspondence of the deterministic KJMA and mKJMA equation with the deterministic MBBG equation is demonstrated theoretically, aspects that allow establishing explicit relationship among the parameters of the deterministic KJMA equation with the deterministic MBBG equation, and those of the deterministic mKJMA equation with the deterministic MBBG equation [5].

The deterministic MBBG equation may to describe unperturbed TGK from the tumor size that is observed and palpated for the first time, named *V*_*obs*_. *V*_*obs*_ is reached for the tumor latency time, which is characteristic when it growths in a type of host. *V*_*obs*_ is less than the tumor size selected by the researcher in *in vitro* and preclinical studies, named *V*_0_. In clinics, *V*_0_ is the tumor size first diagnosed by the doctor [4]. Furthermore, the deterministic MBBG equation contains other parameters, as apoptosis rate; intrinsic growth rate due to mitosis rate, named *α*; and the growth deceleration factor *β*. The parameter *β* is related to the endogenous antiangiogenic process [3–7].

From a mathematical point of view, the deterministic CG equation establishes that the infinite tumor volume named *V*_∞_ (*V*_∞_ is reached when *t* tends to infinite), depends on *V*_0_ for *α* and *β* fixed. From a biophysical point of view, this means that *V*_∞_ depends on the time at which *V*_0_ is observed, in contrast with the natural history of a tumor growing in a given host. Nevertheless, the deterministic MBBG equation establishes that *V*_∞_ depends on *V*_*obs*_ for the same organism carrying a tumor histological variety [4]. The latter is verified in the experiment because the times for which the solid tumor reaches *V*_0_ and *V*_∞_ are well estimated, once *V*_*obs*_ has been observed. This assumes that unperturbed TGK in the selected host is known *a priori*, as in the experiment, in which *V*_∞_ means that the tumor size does not exceed 10% of body weight (ethical standards established for laboratory animals) [3, 4, 6, 7].

The behaviors of untreated TGK that predict deterministic models may be affected by unpredictable fluctuations of their parameters due to endogenous randomness inherent in cancer [8]. Therefore, a diversity of stochastic models is reported in the literature to describe unperturbed TGK [9–12]. Specifically, the stochastic Gompertzian model is used by several authors [13–15].

For unperturbed TGK, the nonlinear Gompertzian stochastic model reported by Ferrante et al. [15] consists in a stochastic differential equation with the initial condition *V* (*t* = 0) = *V*_0_ and assumption *θ*(*t*) = *α* + *ση*(*t*), where *t, V*_0_, *V* (*t*), *θ*(*t*), 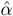, *σ* and *η*(*t*) are the time, initial tumor volume, tumor volume at any instant of time, intrinsic growth rate that varies in time, constant mean value of *θ*(*t*), diffusion coefficient and Gaussian white noise process, respectively. They analytically obtain the maximum likelihood estimators for *α*, named 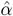; and *β*, named 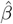 as well as, a discrete-time approximation of the estimator *α*, named 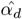, using the Ito formula and trapezoidal rule.

We are not aware that the stochastic version of MBBG equation is reported in the literature. Furthermore, the analytical expressions of 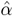 and 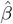 for the stochastic MBBG equation, as well as their respective discrete time approximate estimators are not known. Therefore, the aim of this study is to calculate the parameter estimations in a nonlinear MBBG stochastic model to describe unperturbed TGK. For this, we follow the same ideas of Ferrante et al. [15]. The maximum likelihood estimators and discrete time approximate estimators for intrinsic growth rate and the growth deceleration factor are analytically calculated. The results of this study are discussed from mathematical and biophysical points of view.

## Methods

### Deterministic MBBG model

The ordinary differential equation of the deterministic MBBG model was given by [4]

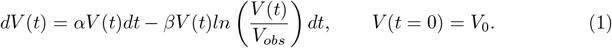

The solution of Equation (1) was

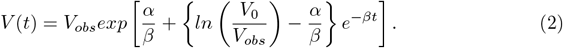

The Equation (2) was named the deterministic MBBG model. From the simulation of this equation, it could be verified easily that the unperturbed TGK was faster when the value of *β* was much smaller than that *α*. This TGK was slower when *β* tends to *α* (*α > β*). If the value of *β* increased with respect to that of *α, V* (*t*) increased over time and tended to zero for *β* much greater than that *α*. This may suggested the existence of a limit condition for *β*, named *β*_*lim*_, which was computed analytically if the square brackets in Equation (2) was zero, resulting

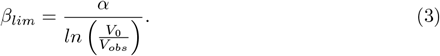

### Stochastic MBBG model

The new stochastic MBBG model was analytically obtained from the deterministic MBBG model [4], following the same mathematical procedure reported by Ferrante et al. [15]. For this, we assumed that *β* remained unchanged, whereas the variability in chemical [16], electrical [17] and mechanical [18] environments of the tumor-induced fluctuations in *α*. Consequently, *α* varied over time. Ferrante et al. [15] proposed a specific behavior for the temporal change of *α*, gave by

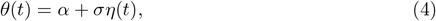

where *α* represented the constant average value of *θ*(*t*), *σ >* 0 was the average diffusion coefficient, and *η*(*t*) was a Gaussian while noise process.

The diffusion coefficient in the solid tumor was considered a tensor due to its anisotropy both in experimental [19] and theoretical [20] studies; nevertheless, in these these two studies and others used the average value of the diffusion coefficient tensor because the mathematical procedure with tensors was cumbersome. This and the objective of this study were why we used the average diffusion coefficient.

The mentioned above allowed that Equation (1) was became to the following stochastic MBBG differential equation, which was given by:

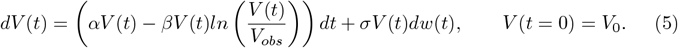

where *W* = {*w*(*t*) : *t* ∈ [0, *T* ]} was the standard Wiener process [15].

### The analytical solution of the stochastic MBBG equation

The Equation (5) was written as follows:

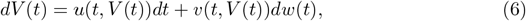

where

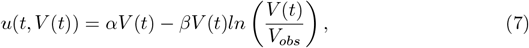

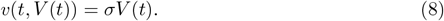

Let 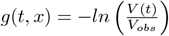, and using the Ito formula (Theorem 4.1.2) reported in [21], we obtained

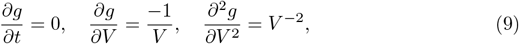

then,

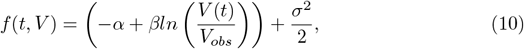

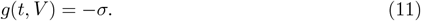

we obtained

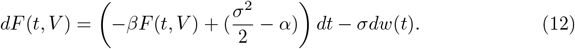

So, we used the **Theorem 2.3.6** reported in [22] to find the solution of *F* (*t, V*), taken *t*_0_ = 0 and calculated the solution of ordinary differential equation *dF* (*t*) = *A*(*t*)*F* (*t*), named Φ(*t*), to solve the Equation (12). Φ(*t*) was given by

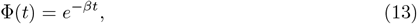

then the solution of Equation (12) was given by

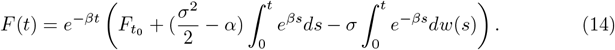

From Equation (14), it could be easily verifed that

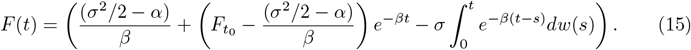

Taken into account that 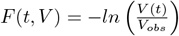 and the initial condition *V* (*t* = 0) = *V*_0_, the final expression for *V* (*t*) was given by

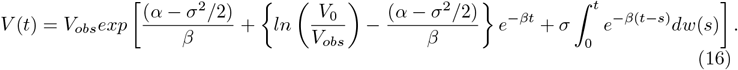

The Equation (16) was the analytic solution for the stochastic MBBG model to describe unperturbed TGK.

### Analysis of stochastic of *V*_∞_

From the analytic solution (16), *V* (*t*) → *V*_∞_, as *t* tended to infinity (*t* → ∞) was given by

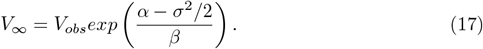

The expression of *V*_∞_ coincided with that of deterministic MBBG model when *σ* = 0.

### Limit conditions for *β* and *σ* in the stochastic MBBG equation

The *β*_*lim*_ for the stochastic MBBG equation (Equation (18)) was analytically computed when the difference of the two terms contained in the square brackets in Equation (16) was equated to zero.

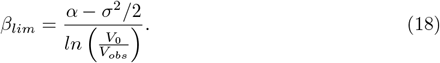

Although in the literature, a limit condition for *σ* (*σ*_*lim*_) had not been documented, for the case of the stochastic MBBG equation, (Equation (19)), it was analytically computed following the same procedure for *β*_*lim*_.

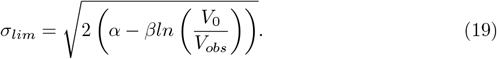

### The maximum likelihood estimator of *α* and *β* for stochastic MBBG equation

The probability density (Radon-Nikodym derivative), named *L*(*a, V*), was calculated using the Theorem 3 reported in [23] and making the following substitutions in Equation (1)

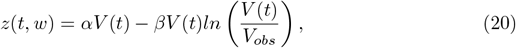

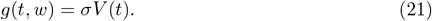

The expression of *L*(*a, V*) was given by

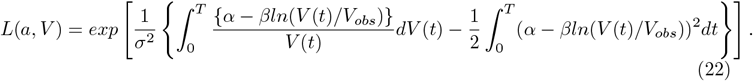

### The maximum likelihood estimator of *α*

The maximum likelihood estimator of *α* 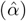 was obtained from the partial derivate of Equation(22) with respect to *α* and setting it to zero. Consequently, we obtained

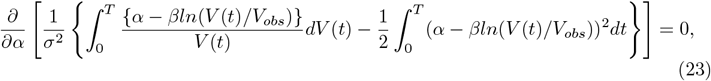

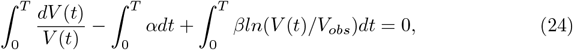

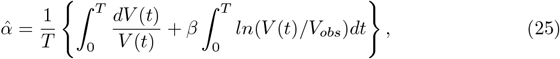

The Equation (25) was rewritten as

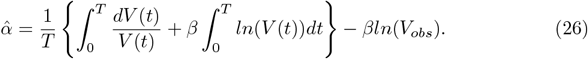

The estimator 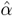 in Equation (26) differed from that for stochastic CG equation [15] in the term −*βln*(*V*_*obs*_).

### The maximum likelihood estimator of *β*

The maximum likelihood estimator of *β* 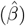 was obtained analytically following the same mathematical procedure for 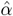. In this case, we take the partial derivate of Equation (22) with respect to *β*, yielding

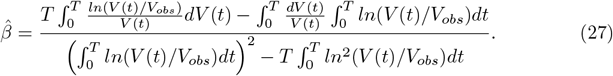

If we expanded the logaritms in Equation (27), we obtained

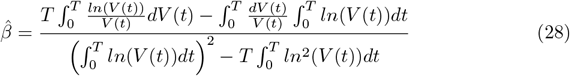

The estimator 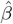 in Equation (28) exactly coincided with that for stochastic CG equation reported in [15].

### Discrete time approximation

#### Discrete time approximation the estimator 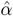

The discrete-time approximation of the estimator 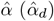 was obtained from Equation (26) and using the rule of 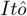 (Appendix I [21]), we obtained

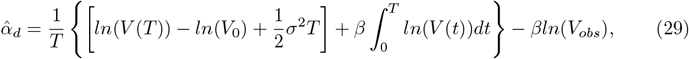

then

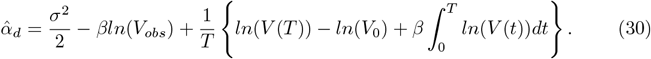

The rule of trapezoid was used to solve approximately the integral in Equation (30), yielding

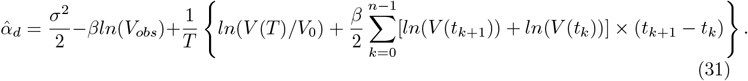

where *n* was the total number of discrete data of *V* (*t*). We used *n* = 128, as suggested in [15].

#### Discrete time approximation the estimator 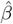

The discrete-time approximation the estimator 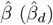 was obtained from Equation (37) and using the rule of 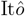 (Appendix I [21]), we obtained

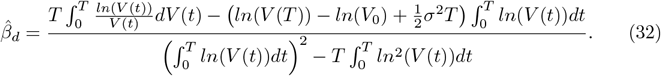

We used again the rule of 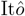 (Appendix I [21]) to change the integration variable *dV* (*t*) into *dt* in Equation (32). For this, 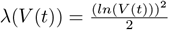, then 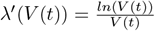 and 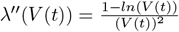, yielding

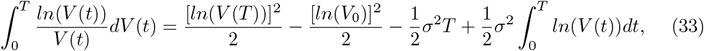

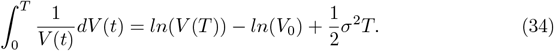

Substituting Equations (33) and (34) in Equation (32), resulted the following equation for 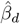

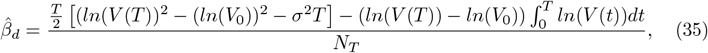

where *N*_*T*_ was given by

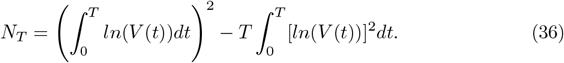

The estimator for 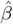 for a set of *n* discrete data of *V* (*t*) was calculated using the rule of trapezoid.

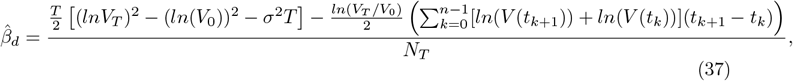

with

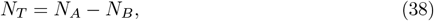

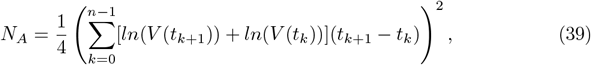

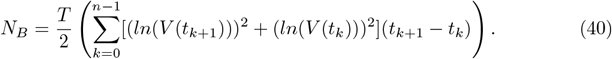

## Simulation

The plots of *V* (*t*) (in *cm*^3^) versus *t* (in *days*) for the deterministic and stochastic MBBG equations were shown. For the case of the deterministic MBBG equation, different values of *β* (0.06, 0.17, 0.22, 0.34, 0.50 and 1.00 *days*^−1^) and *V*_*obs*_ (0.005, 0.050, 0.100 and 0.500 *cm*^3^) were used, keeping constant the values of *V*_0_ (0.5 *cm*^3^), *α* (0.8 *days*^−1^) and *σ* (0.000). For the case of the stochastic MBBG equation, we used these same values of *β, V*_*obs*_, *V*_0_, *α* and different values of *σ* (0.025, 0.100, 0.500, 0.800, 1.000 and 1.200). For the stochastic MBBG equation, the mean, standard deviation, median, and quantile of 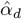 were calculated. For this, we used *T* = 30 *days*; the same values of *α, V*_0_, *β* and *σ*; and 500 simulations were conducted for each experiment. The value of *T* was fixed from the experiment [4]. The number of partitions for each simulation overtime was *n* = 128. Ferrante et al. [15] demostrated that the difference among means of 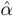 for *n* = 128, 256, and 512 was ≤ 0.002 *days*^−1^, finding obeserved for their four cases *SIM* 1, *SIM* 2, *SIM* 3 and *SIM* 4 (see Table 1 of [15]). This maximum difference for 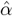 was not relevant in the experiment [3, 4, 7, 8]. This was why, we fixed *n* = 128 in this study.

**Table 1.** Values of *V*_∞_ for different values of *σ* and *V*_*obs*_. *V*_∞_, *V*_*obs*_ and *σ* represented the infinite tumor volume, tumor size observed and reported for the first time and average diffusion coefficient, respectively.

| $V_{obs} \text{ (cm}^3\text{)}$ | $V_\infty \text{ (cm}^3\text{)}$ | | | | | | |
| --- | --- | --- | --- | --- | --- | --- | --- |
| | $\sigma = 0.000$ | $\sigma = 0.025$ | $\sigma = 0.100$ | $\sigma = 0.500$ | $\sigma = 0.800$ | $\sigma = 1.000$ | $\sigma = 1.200$ |
| 0.005 | 0.5529 | 0.5519 | 0.5369 | 0.2650 | 0.0841 | 0.0291 | 0.0080 |
| 0.050 | 5.5297 | 5.5196 | 5.3695 | 2.6507 | 0.8418 | 0.2919 | 0.0800 |
| 0.100 | 11.0595 | 11.0399 | 10.7390 | 5.3015 | 1.6836 | 0.5839 | 0.1600 |
| 0.500 | 55.2979 | 55.1963 | 53.6951 | 26.5078 | 8.4180 | 2.9199 | 0.8004 |

## Results

The *V* (*t*) versus t plot for the deterministic MBBG equation showed in Figure 1, whereas this plot for the stochastic MBBG equation was displayed in Figures 2-4. The *V* (*t*) versus *t* plot for the deterministic MBBG equation was showed for *V*_*obs*_ = 0.005 *cm*^3^ (Figure 1A), *V*_*obs*_ = 0.050 *cm*^3^ (Figure 1B), *V*_*obs*_ = 0.100 *cm*^3^ (Figure 1C) and *V*_*obs*_ = 0.500 *cm*^3^ (Figure 1D), different values of *β*, and values of *α, V*_0_ and *σ* were kept constants. This figure evidenced that *V* (*t*) grew rapidly for the lowest values of *β* and all values of *V*_*obs*_. Nevertheless, *V* (*t*) remained constant (*β* = 0.17 *days*^−1^) or decreased to a non-zero value (*β* = 0.22 *days*^−1^) or zero (other values of *β*) for the lowest value of *V*_*obs*_. These behaviors of *V* (*t*) changed markedly when *V*_*obs*_ increased (*V*_*obs*_ tended to *V*_0_) for each *β* value. In this case, TGK was fast for the highest values of *V*_*obs*_.

**Fig 1.**
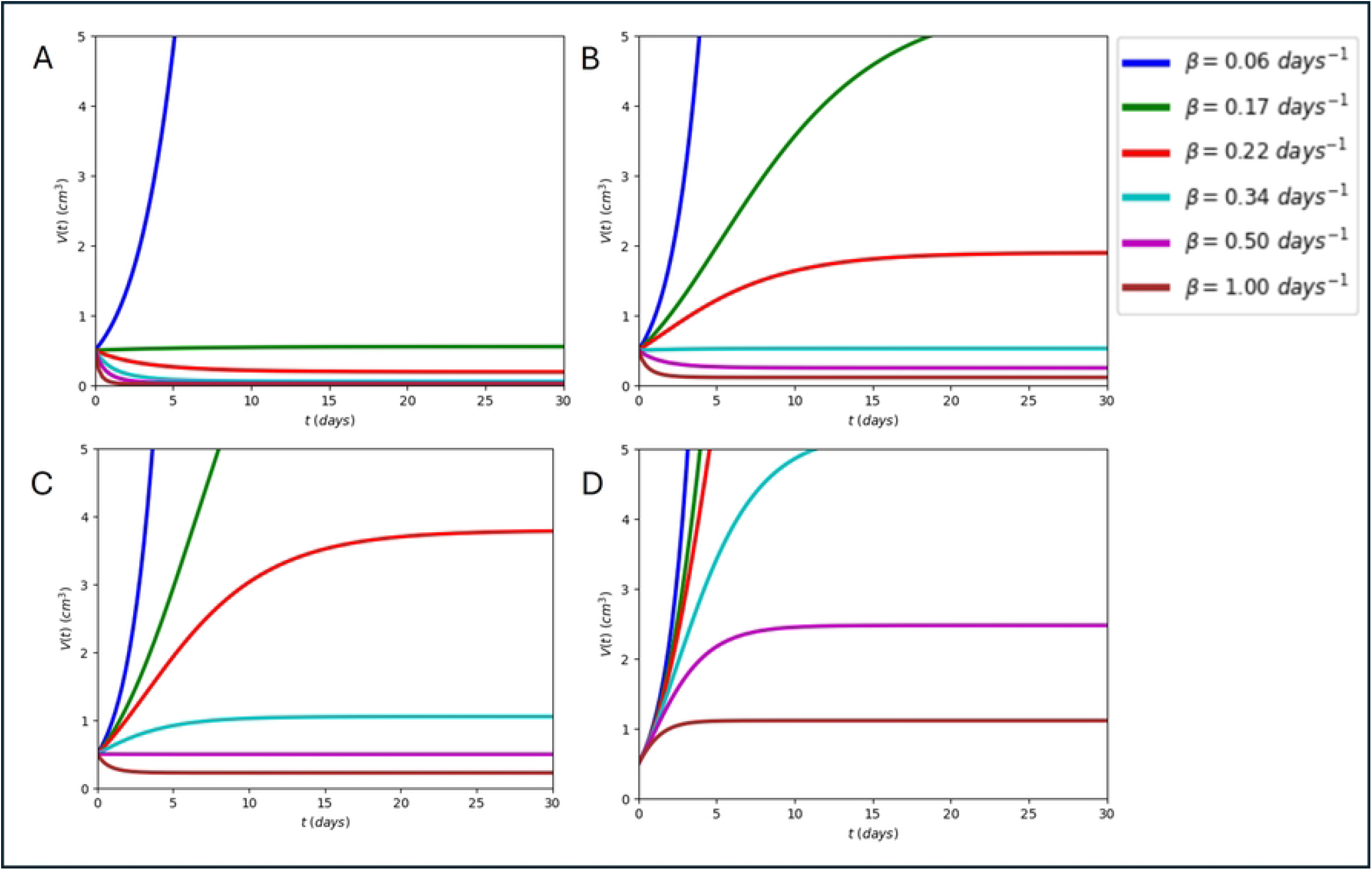
Plot of *V* (*t*) versus *t*. Plot obtained with deterministic MBBG equation for *V*_0_ = 0.5 *cm*^3^, *α* = 0.8 *days*^−1^, *σ* = 0.000 and different values of *β*. A) *V*_*obs*_ = 0.005 *cm*^3^, B) *V*_*obs*_ = 0.050 *cm*^3^, C) *V*_*obs*_ = 0.100 *cm*^3^, D) *V*_*obs*_ = 0.500 *cm*^3^.

**Fig 2.**
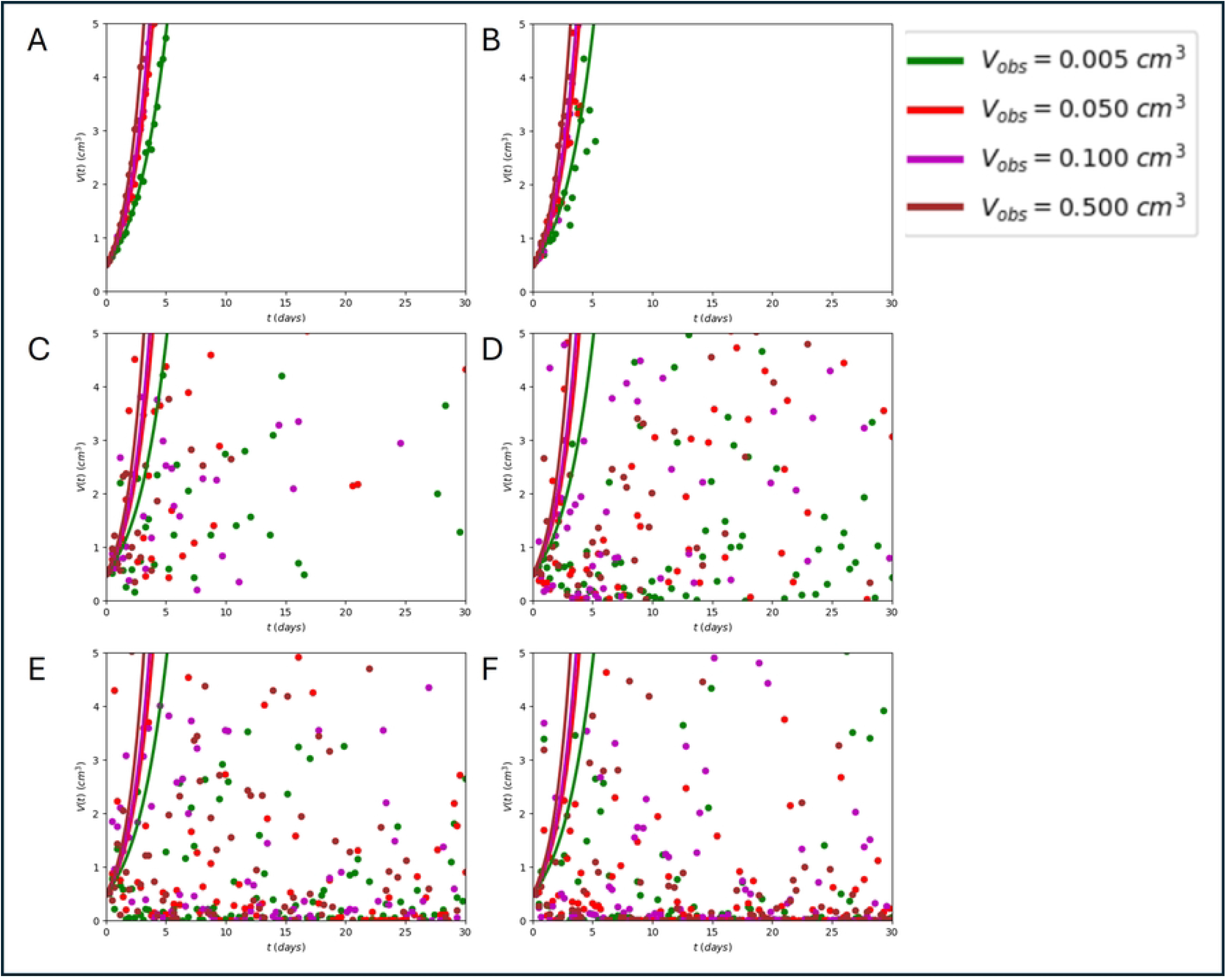
Plot of *V* (*t*) versus *t* for the stochastic MBBG equation. Plot obtained for *α* = 0.8 *days*^−1^, *β* = 0.06 *days*^−1^, *V*_0_ = 0.5 *cm*^3^, and different values of *V*_*obs*_. **A)** *σ* = 0.025, **B)** *σ* = 0.100, **C)** *σ* = 0.500, **D)** *σ* = 0.800, **E)** *σ* = 1.000, **F)** *σ* = 1.200.

The *V* (*t*) versus *t* plots for the stochastic MBBG equation were shown for constant values of *α, β* and *V*_0_; different values of *V*_*obs*_ for *σ* = 0.025 (Figures 2-4A), *σ* = 0.100 (Figure 2-4B), *σ* = 0.500 (Figure 2-4C), *σ* = 0.800 (Figure 2-4D), *σ* = 1.000 (Figure 2-4E) and *σ* = 1.200 (Figure 2-4F). Figures 2, 3, and 4 showed the results for *β* values of 0.06, 0.2, and 0.4 *days*^−1^, respectively.

**Fig 3.**
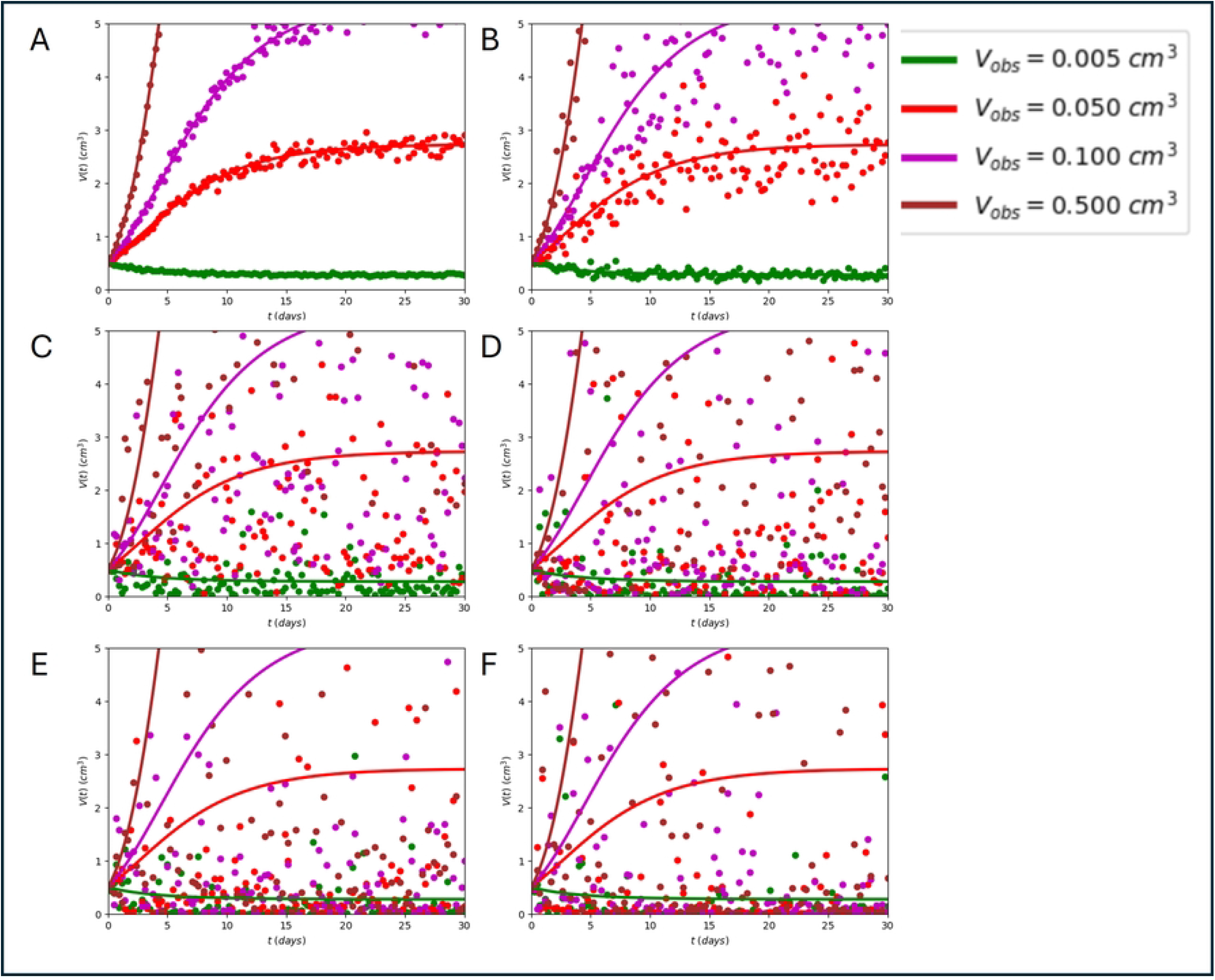
Plot of *V* (*t*) versus *t* for the stochastic MBBG equation. Plot obtained for *α* = 0.8 *days*^−1^, *β* = 0.2 *days*^−1^, *V*_0_ = 0.5 *cm*^3^, and different values of *V*_*obs*_. **A)** *σ* = 0.025, **B)** *σ* = 0.100, **C)** *σ* = 0.500, **D)** *σ* = 0.800, **E)** *σ* = 1.000, **F)** *σ* = 1.200.

**Fig 4.**
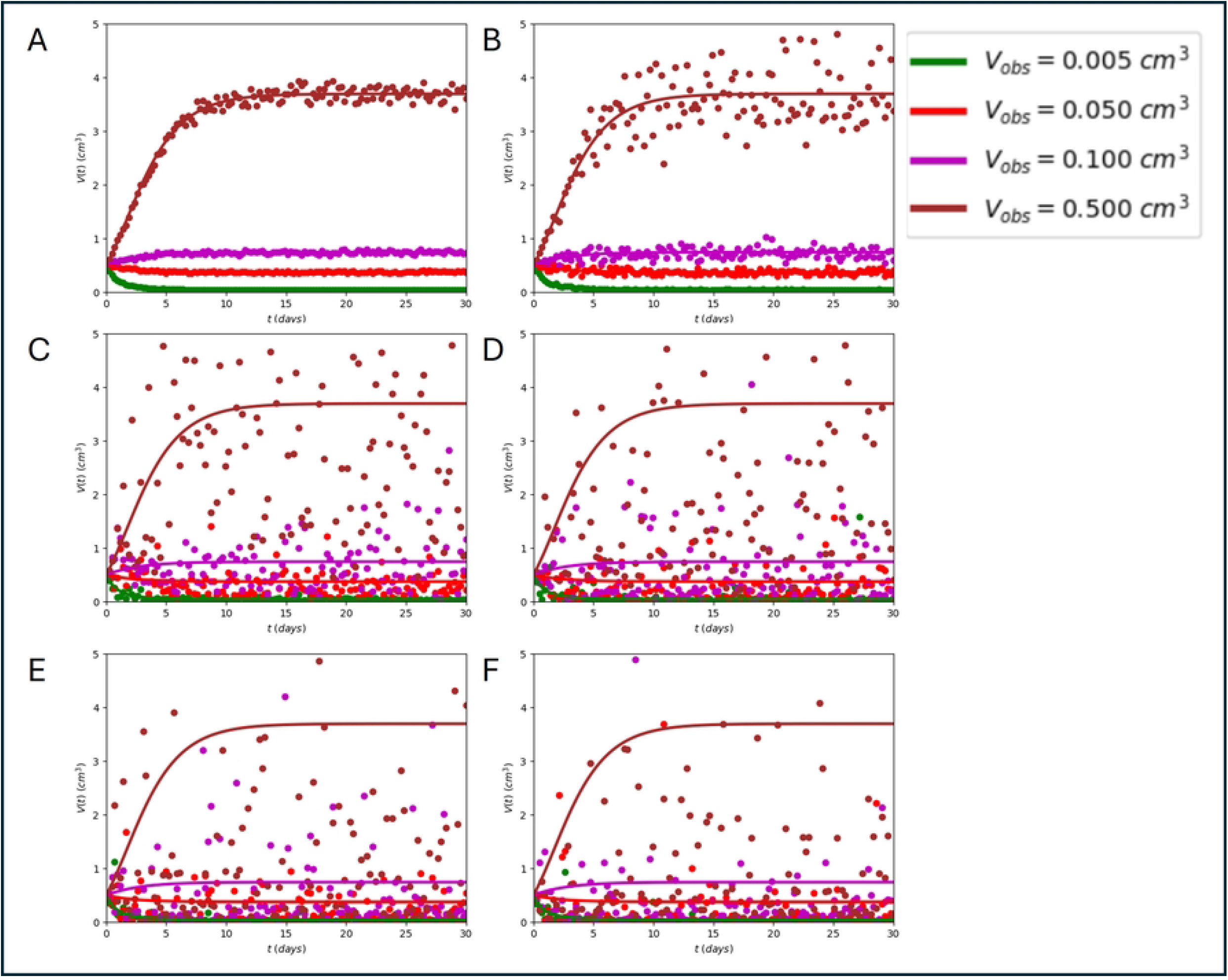
Plot of *V* (*t*) versus *t* for the stochastic MBBG equation. Plot obtained for *α* = 0.8 *days*^−1^, *β* = 0.4 *days*^−1^, *V*_0_ = 0.5 *cm*^3^, and different values of *V*_*obs*_. **A)** *σ* = 0.025, **B)** *σ* = 0.100, **C)** *σ* = 0.500, **D)** *σ* = 0.800, **E)** *σ* = 1.000, **F)** *σ* = 1.200.

Figures 2-4 evidenced that the results obtained with deterministic (solid lines) and stochastic (points) MBBG equations agreed for lowest *σ* values (0.000 ≤ *σ* ≤ 0.025) and all values of *β* and *V*_*obs*_. This finding was observed for 0.025 *< σ* ≤ 0.100, the lowest *β* value, and all values of *V*_*obs*_. Nevertheless, a greater degree of stochasticity (greater dispersion of tumor volume) was predicted by the stochastic MBBG equation for ≤ 0.500 ≤ *σ* ≤ 0.800 and each *β* value (Figures 2-4C,D), being marked for *β* ≤ 0.2 days-1. This degree of stochasticity began to decrease from *σ >* 0.800 for each *β* value and all values of *V*_*obs*_, being marked for the highest values of *σ* and *β*. Furthermore, Figures 2-4 exhibited that the highest degree of stochasticity was observed for *β* = 0.2 *days*^−1^.

The Figure 1 and the dependence of the degree of stochasticity on the values of *β* and *σ* shown in Figures 2-4 suggested the existence of limit/threshold values for *β* and *σ*, which were calculated analytically from Equation (18) for *β*_*lim*_ and Equation (19) for *σ*_*lim*_, and displayed graphically in Figure 5. The Figure 5 showed the plots *σ* versus *β* (Figure 5A) and *ln*(*σ*) versus *ln*(*β*) (Figure 5B) obtained with the stochastic MBBG equation for four values of *V*_*obs*_ (0.005, 0.050, 0.100 and 0.500 *cm*^3^) and values of *α* and *V*_0_ kept constants. The results of this figure revealed that *β*_*lim*_ was undefined and *σ*_*lim*_ = 1.264 when *V*_*obs*_ = *V*_0_. That was why, *V*_*obs*_ = 0.5 *cm*^3^ was not used for simulations in Figure 5.

**Fig 5.**
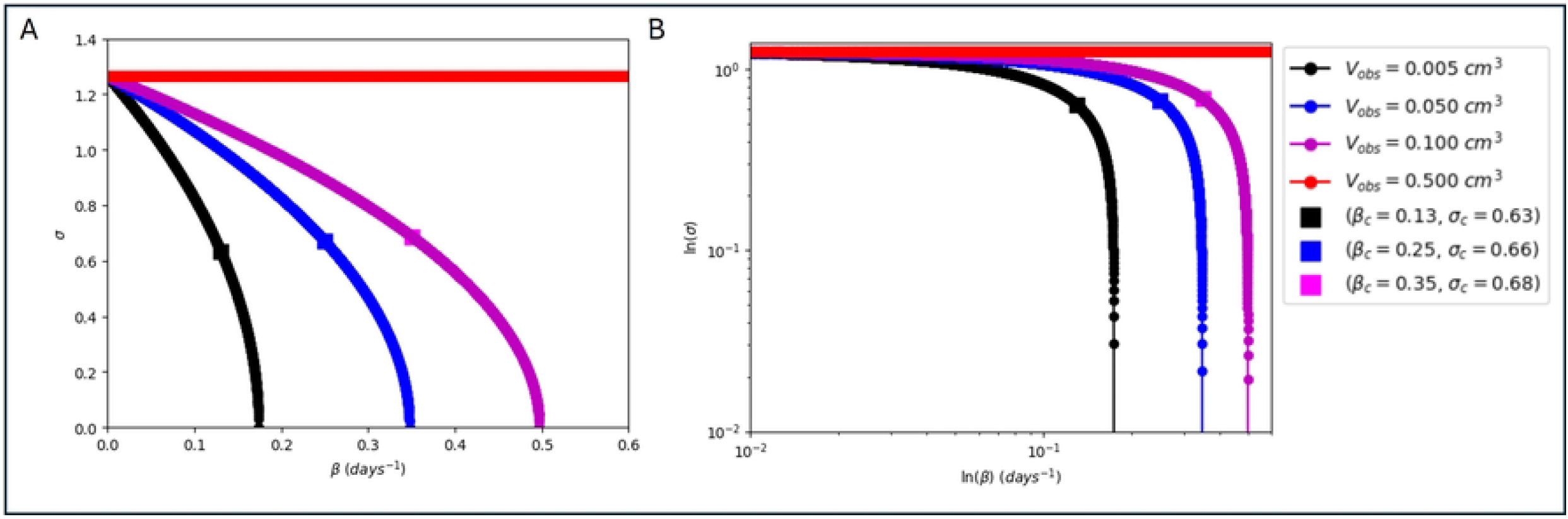
Graphic representation of the existence of limit values of *β* (*β*_*lim*_) and *σ* (*σ*_*lim*_). **A) Plot of** *σ* **versus** *β*. **B) Plot of** *ln*(*σ*) **versus** *ln*(*β*). **Both subplots were made for** *α* = 0.8 *days*^−1^, *V*_0_ = 0.5 *cm*^3^ **and four values of** *V*_*obs*_.

For each value of *V*_*obs*_, *σ* decreased nonlinearly from *σ*_*lim*_ up to zero when *β* increased, being marked for the lowest value of *V*_*obs*_. This nonlinear decrease in *σ* had an abrupt change in slope at the point (*β*_*c*_, *σ*_*c*_), which was indicated with the black square on each curve corresponding to *V*_*obs*_ (Figure 5). This change in slope defined a first slow part (from *σ*_*lim*_ up to *σ*_*c*_) and another very fast one (from *σ*_*c*_ up to *σ* = 0.000). The point (*β*_*c*_, *σ*_*c*_) indicated the coordinates of the black square showed in Figure 5.

It was interesting that *σ* = 0.000 was observed for *β* = 0.1737 *days*^−1^ (*V*_*obs*_ = 0.025 *cm*^3^), *β* = 0.3474 *days*^−1^ (*V*_*obs*_ = 0.050 *cm*^3^), and *β* = 0.4970 *days*^−1^ (*V*_*obs*_ = 0.100 *cm*^3^). In these cases, the deterministic and stochastic MBBG equations coincided. The zone enclosed between the x-axis (*β* values), the y-axis (*σ* values), and the curve *σ* versus *β* for each value of *V*_*obs*_, defined the set of allowable *β* and *σ* values (parameter portrait) for which the unperturbed solid tumor growth only existed and grew (active and functional cancer). The untreated solid tumor did not exist for the other values of *β* and *σ* outside of this parameter portrait (zone to the right of each curve corresponding to *V*_*obs*_).

The Figure 6 showed the behaviors of 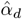 versus 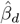 for different values of *σ* (0.000, 0.025, 0.100, 0.500, 0.800, 1.000 and 1.200) and *β* (from 0.01 to 0.6 *days*^−1^). These behaviors were exhibited for *V*_*obs*_ = 0.005 *cm*^3^ (Figure 6A), 0.05 *cm*^3^ (Figure 6B), 0.1 *cm*^3^ (Figure 6C) and 0.5 *cm*^3^ (Figure 6D). The Figure 6 was obtained with Equation (31) (for 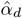 and Equation (37) (for 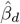 and revealed that 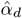 did not change with the increase of 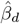 for the case *σ* = 0.000. For *σ* ≠ 0, a nonlinear behavior with a peak (minimum value of 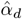) was observed in the 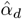 versus 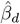 parameter portrait. This nonlinear behaviour of 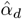 versus 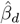, and the positioning and morphology of the peak depended essentially on *σ* and *V*_*obs*_.

**Fig 6.**
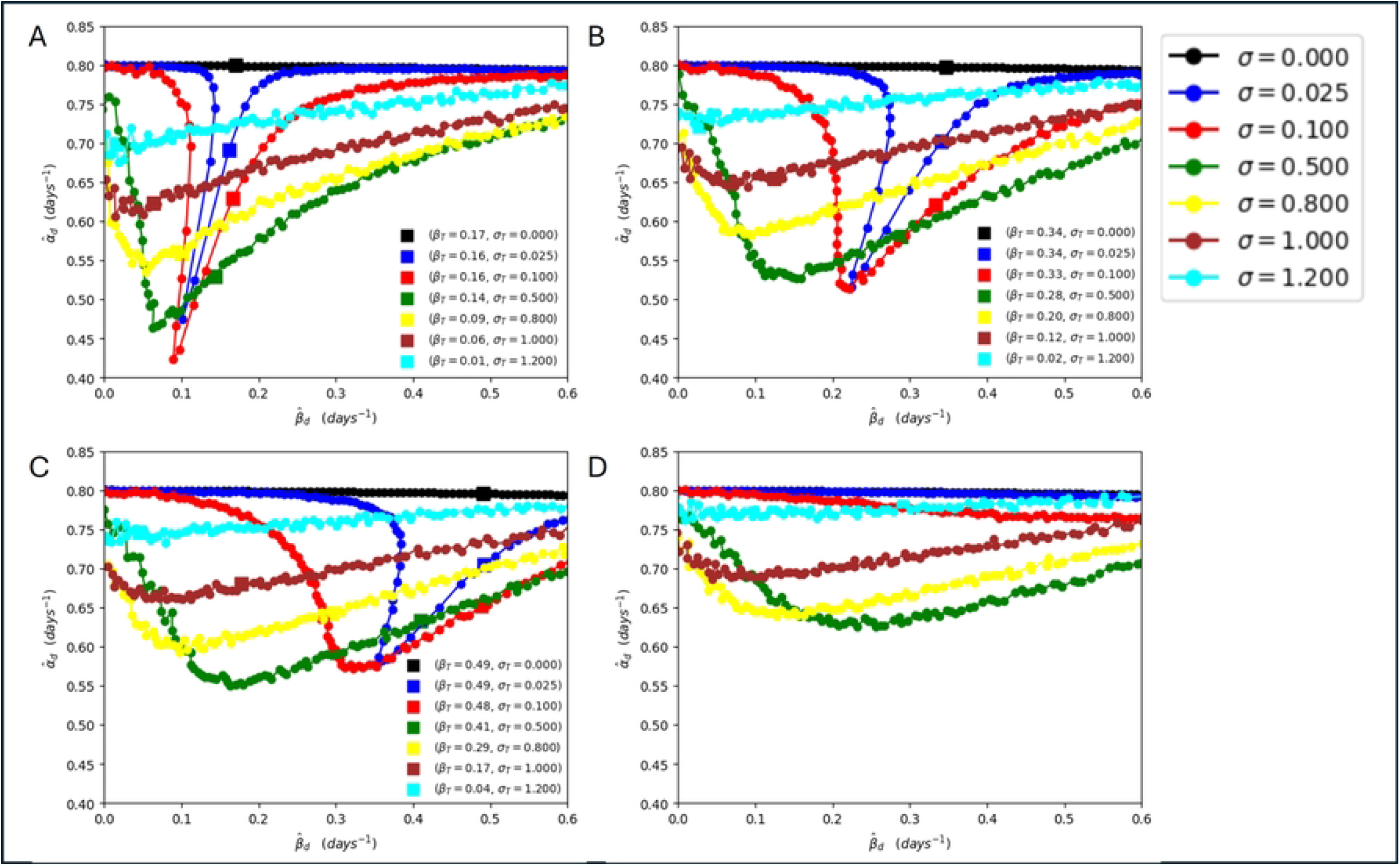
Plot of 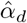 versus 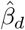. **Plot obtained for** *α* **and** *V*_0_ **constants, seven values of** *σ*, 0.01≤ *β* ≤ 0.6 *day*^−1^, **and four values of** *V*_*obs*_**: A)** *V*_*obs*_ = 0.005 *cm*^3^ **B)** *V*_*obs*_ = 0.05 *cm*^3^ **C)** *V*_*obs*_ = 0.1 *cm*^3^ **D)** *V*_*obs*_ = 0.5 *cm*^3^.

The sharpest peak (narrowest and greatest amplitude) corresponded for *σ* = 0.025 and *V*_*obs*_ = 0.005 *cm*^3^, and its minimum value of 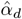, was observed for 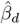 close to 0.1 *days*^−1^ (Figure 6A). Furthermore, this figure showed another peak of amplitude similar to that corresponding to *σ* = 0.025 and *V*_*obs*_ = 0.005 *cm*^3^ but wider for *σ* = 0.100, *V*_*obs*_ = 0.005 *cm*^3^ and the same values of 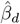 and *V*_*obs*_. The peaks of 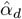 were lower in amplitude, wider and minimum values of 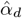 shifted more to the left (respect to 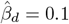 *days*^−1^) when *σ* increased for *V*_*obs*_ = 0.005 *cm*^3^. Nevertheless, the amplitudes, width, and positioning of all peaks shown in Figure 6A were smaller, larger, and shifted more to right (relative to 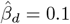 *days*^−1^) when *V*_*obs*_ increased for each *σ* value. The left (active tumor) and right (inactive tumor) branches of 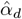 versus 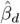 represented the growth and decrease of TGK, respectively. These two branches were separated by the point (*β*_*T*_, *σ*_*T*_) for each curve that corresponded to the values of *σ* and *V*_*obs*_, except in *V*_*obs*_ = 0.5 *cm*^3^, for which the square was not observed in any of the curves for all sigma values.

The Table 1 showed *V*_∞_ values calculated from the stochastic MBBG equation for *α* = 0.8 (*days*^−1^); *β* = 0.17 (*days*^−1^); and different values of *V*_*obs*_ (0.005, 0.050, 0.100 and 0.500 *cm*^3^) and *σ* (0.000, 0.025, 0.100, 0.500, 0.800, 1.000 and 1.200). This table revealed that *V*_∞_ decreased when *σ* increased for each constant value of *V*_*obs*_.

The Tables 2-5 showed the values of parameters obtained from descriptive statistic to estimate mean of 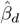, mean of 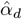, standard deviation of 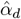, median of 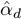 and quartile of 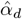 for three values of *σ* (0.500, 0.800 and 1.200), *V*_*obs*_ = 0.5 *cm*^3^, *n* = 128, *T* = 30 *days*, number of simulation (discrete trajectories) 500, *β* (from 0.01 to 0.7 *days*^−1^), and *V*_*obs*_ values of 0.005 *cm*^3^ (Table 2), 0.005 *cm*^3^ (Table 3), 0.1 *cm*^3^ (Table 4) and 0.05 *cm*^3^ (Table 5). The results of Tables 2-5— evidenced that the mean of 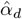 depended on *σ, β* and *V*_*obs*_.

**Table 2.**
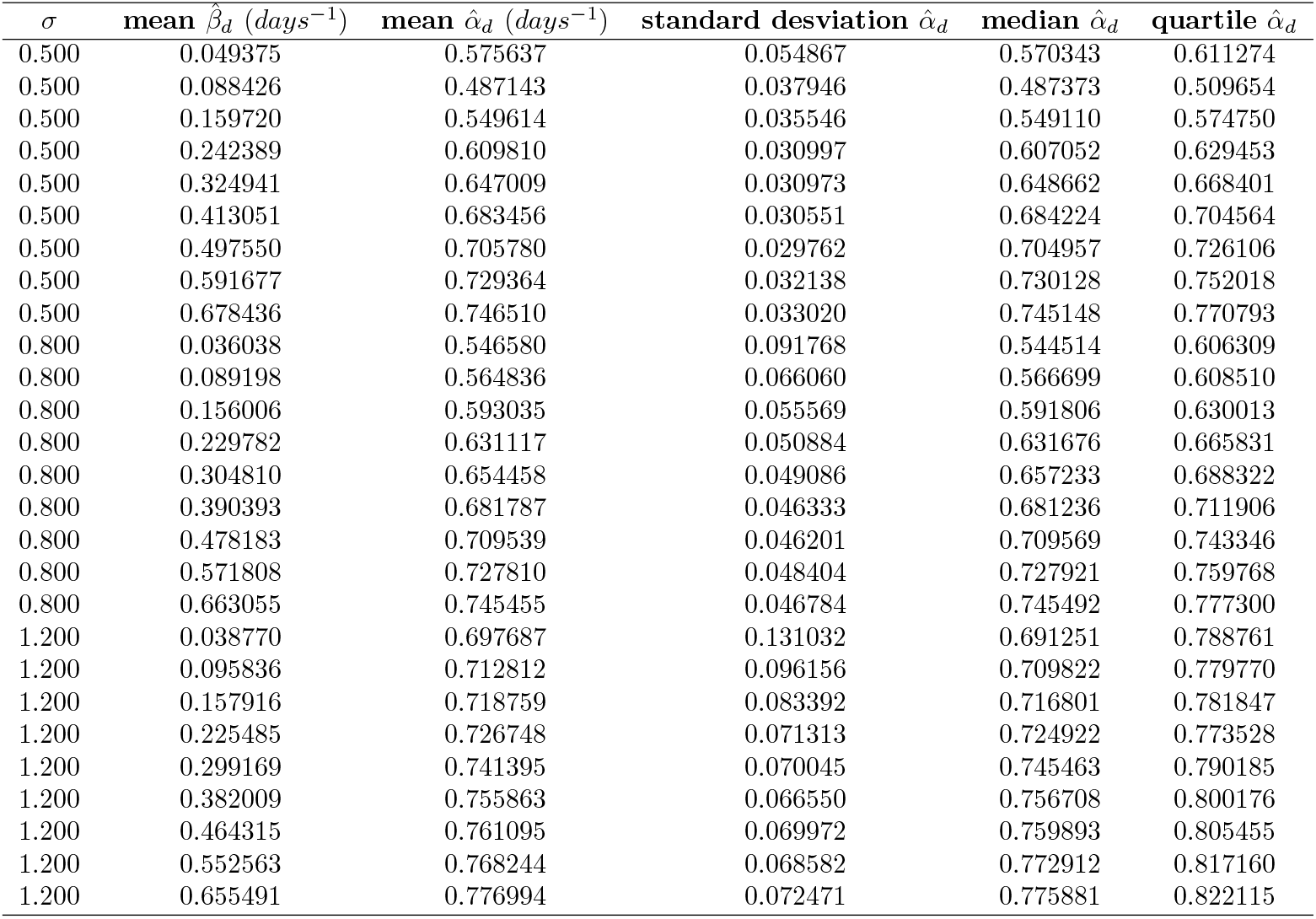
Descriptive statistic for *V*_*obs*_ = 0.005 *cm*^3^.

**Table 3.** Descriptive statistic for *V*_*obs*_ = 0.05 *cm*^3^.

| $\sigma$ | mean $\hat{\beta}_d \text{ (days}^{-1}\text{)}$ | mean $\hat{\alpha}_d \text{ (days}^{-1}\text{)}$ | standard desviation $\hat{\alpha}_d$ | median $\hat{\alpha}_d$ | quartile $\hat{\alpha}_d$ |
| --- | --- | --- | --- | --- | --- |
| 0.500 | 0.059094 | 0.663125 | 0.055503 | 0.665829 | 0.702956 |
| 0.500 | 0.097647 | 0.558880 | 0.039628 | 0.557692 | 0.585678 |
| 0.500 | 0.143941 | 0.530579 | 0.033475 | 0.529519 | 0.553501 |
| 0.500 | 0.205471 | 0.548556 | 0.029658 | 0.551203 | 0.567713 |
| 0.500 | 0.278116 | 0.582605 | 0.029218 | 0.582385 | 0.602429 |
| 0.500 | 0.358967 | 0.615294 | 0.029294 | 0.614943 | 0.635204 |
| 0.500 | 0.442532 | 0.644400 | 0.027506 | 0.642734 | 0.662397 |
| 0.500 | 0.542683 | 0.684740 | 0.027827 | 0.685074 | 0.702310 |
| 0.500 | 0.641505 | 0.713679 | 0.030268 | 0.713885 | 0.734969 |
| 0.800 | 0.041037 | 0.619489 | 0.089024 | 0.616832 | 0.679676 |
| 0.800 | 0.085430 | 0.584963 | 0.068416 | 0.585371 | 0.630210 |
| 0.800 | 0.141024 | 0.595628 | 0.054280 | 0.597855 | 0.630388 |
| 0.800 | 0.211744 | 0.621048 | 0.045606 | 0.618015 | 0.649020 |
| 0.800 | 0.279105 | 0.636198 | 0.045213 | 0.635952 | 0.669290 |
| 0.800 | 0.357673 | 0.660172 | 0.045014 | 0.660156 | 0.689675 |
| 0.800 | 0.441363 | 0.682393 | 0.045823 | 0.684211 | 0.715877 |
| 0.800 | 0.534073 | 0.705930 | 0.047257 | 0.703237 | 0.737815 |
| 0.800 | 0.639265 | 0.732751 | 0.048094 | 0.734873 | 0.765255 |
| 1.200 | 0.036631 | 0.731912 | 0.133870 | 0.734715 | 0.822078 |
| 1.200 | 0.087051 | 0.739848 | 0.100492 | 0.741611 | 0.804387 |
| 1.200 | 0.145669 | 0.742644 | 0.076873 | 0.740924 | 0.792055 |
| 1.200 | 0.209612 | 0.746046 | 0.074125 | 0.743630 | 0.797176 |
| 1.200 | 0.280440 | 0.758308 | 0.072037 | 0.759477 | 0.805309 |
| 1.200 | 0.357849 | 0.764483 | 0.071000 | 0.767002 | 0.810065 |
| 1.200 | 0.439900 | 0.764193 | 0.067075 | 0.767822 | 0.807775 |
| 1.200 | 0.538450 | 0.775503 | 0.067648 | 0.778491 | 0.823160 |
| 1.200 | 0.637107 | 0.779338 | 0.072407 | 0.780292 | 0.832488 |

**Table 4.** Descriptive statistic for *V*_*obs*_ = 0.1 *cm*^3^.

| $\sigma$ | mean $\hat{\beta}_d \text{ (days}^{-1}\text{)}$ | mean $\hat{\alpha}_d \text{ (days}^{-1}\text{)}$ | standard desviation $\hat{\alpha}_d$ | median $\hat{\alpha}_d$ | quartile $\hat{\alpha}_d$ |
| --- | --- | --- | --- | --- | --- |
| 0.500 | 0.060431 | 0.679049 | 0.055262 | 0.679963 | 0.715054 |
| 0.500 | 0.106459 | 0.595051 | 0.040968 | 0.595250 | 0.620879 |
| 0.500 | 0.153051 | 0.563559 | 0.032500 | 0.564233 | 0.584801 |
| 0.500 | 0.208228 | 0.563810 | 0.029581 | 0.561814 | 0.583703 |
| 0.500 | 0.274245 | 0.584351 | 0.028833 | 0.583304 | 0.603252 |
| 0.500 | 0.348331 | 0.606142 | 0.028607 | 0.606320 | 0.625888 |
| 0.500 | 0.435742 | 0.640286 | 0.027815 | 0.640245 | 0.657510 |
| 0.500 | 0.533919 | 0.679926 | 0.027854 | 0.679942 | 0.698753 |
| 0.500 | 0.630627 | 0.705241 | 0.028817 | 0.705595 | 0.724841 |
| 0.800 | 0.041380 | 0.628952 | 0.093342 | 0.634377 | 0.696339 |
| 0.800 | 0.089100 | 0.602102 | 0.065059 | 0.599456 | 0.646738 |
| 0.800 | 0.141999 | 0.604060 | 0.053429 | 0.603120 | 0.638425 |
| 0.800 | 0.202133 | 0.621647 | 0.049907 | 0.623991 | 0.654018 |
| 0.800 | 0.271814 | 0.637580 | 0.046491 | 0.640753 | 0.669527 |
| 0.800 | 0.349913 | 0.657315 | 0.042523 | 0.657886 | 0.684946 |
| 0.800 | 0.436507 | 0.686295 | 0.045127 | 0.685285 | 0.715453 |
| 0.800 | 0.528778 | 0.704907 | 0.049558 | 0.704350 | 0.738063 |
| 0.800 | 0.629533 | 0.731023 | 0.046599 | 0.731311 | 0.763820 |
| 1.200 | 0.034615 | 0.748626 | 0.136641 | 0.757263 | 0.839195 |
| 1.200 | 0.087029 | 0.744565 | 0.097512 | 0.745116 | 0.810354 |
| 1.200 | 0.145179 | 0.748698 | 0.076680 | 0.744891 | 0.793648 |
| 1.200 | 0.207225 | 0.750673 | 0.072641 | 0.749925 | 0.803815 |
| 1.200 | 0.277830 | 0.758749 | 0.067281 | 0.759064 | 0.804568 |
| 1.200 | 0.351509 | 0.766681 | 0.068641 | 0.767715 | 0.809578 |
| 1.200 | 0.435483 | 0.768359 | 0.068186 | 0.770447 | 0.819568 |
| 1.200 | 0.535715 | 0.779170 | 0.067292 | 0.781356 | 0.824062 |
| 1.200 | 0.633957 | 0.787923 | 0.073047 | 0.789560 | 0.833637 |

**Table 5.** Descriptive statistic for *V*_*obs*_ = 0.5 *cm*^3^.

| $\sigma$ | mean $\hat{\beta}_d \text{ (days}^{-1}\text{)}$ | mean $\hat{\alpha}_d \text{ (days}^{-1}\text{)}$ | standard desviation $\hat{\alpha}_d$ | median $\hat{\alpha}_d$ | quartile $\hat{\alpha}_d$ |
| --- | --- | --- | --- | --- | --- |
| 0.500 | 0.066408 | 0.720219 | 0.058383 | 0.717990 | 0.762618 |
| 0.500 | 0.120046 | 0.659426 | 0.043580 | 0.659836 | 0.688933 |
| 0.500 | 0.175963 | 0.638752 | 0.035271 | 0.638318 | 0.664360 |
| 0.500 | 0.232487 | 0.626044 | 0.029094 | 0.626557 | 0.646869 |
| 0.500 | 0.296791 | 0.633601 | 0.029664 | 0.634775 | 0.653949 |
| 0.500 | 0.366099 | 0.646557 | 0.029363 | 0.646228 | 0.667670 |
| 0.500 | 0.449255 | 0.670016 | 0.027698 | 0.670367 | 0.688514 |
| 0.500 | 0.534631 | 0.688310 | 0.028965 | 0.688328 | 0.707913 |
| 0.500 | 0.643635 | 0.723907 | 0.029322 | 0.724011 | 0.742666 |
| 0.800 | 0.048285 | 0.678351 | 0.090057 | 0.674744 | 0.735004 |
| 0.800 | 0.098479 | 0.645883 | 0.063998 | 0.646691 | 0.688604 |
| 0.800 | 0.151272 | 0.644899 | 0.053695 | 0.646070 | 0.681932 |
| 0.800 | 0.210881 | 0.648897 | 0.049214 | 0.649149 | 0.679874 |
| 0.800 | 0.279749 | 0.663532 | 0.045295 | 0.663455 | 0.693093 |
| 0.800 | 0.356520 | 0.680708 | 0.045901 | 0.681533 | 0.713389 |
| 0.800 | 0.431968 | 0.690222 | 0.045675 | 0.689092 | 0.722271 |
| 0.800 | 0.528162 | 0.716110 | 0.047083 | 0.716550 | 0.747381 |
| 0.800 | 0.626854 | 0.734296 | 0.045561 | 0.733710 | 0.766260 |
| 1.200 | 0.036612 | 0.773071 | 0.133989 | 0.773145 | 0.860348 |
| 1.200 | 0.085457 | 0.772755 | 0.094649 | 0.770114 | 0.833528 |
| 1.200 | 0.143107 | 0.776245 | 0.078091 | 0.773285 | 0.829685 |
| 1.200 | 0.203313 | 0.767822 | 0.071699 | 0.762230 | 0.816546 |
| 1.200 | 0.272251 | 0.775338 | 0.069266 | 0.774389 | 0.831057 |
| 1.200 | 0.350044 | 0.779637 | 0.066809 | 0.773838 | 0.821163 |
| 1.200 | 0.435701 | 0.777254 | 0.072544 | 0.779875 | 0.828680 |
| 1.200 | 0.532444 | 0.785316 | 0.070648 | 0.786423 | 0.835037 |
| 1.200 | 0.626697 | 0.792852 | 0.065917 | 0.790148 | 0.835654 |

## Discussion

This study reports for the first time the stochastic version of MBBG equation for the unperturbed TGK, the maximum likelihood estimators for 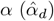 and 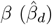, and the discrete time approximations for 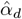 and 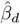 estimators, in terms of *V*_*obs*_, *V*_0_, *σ, α* and *β* (specifically the difference of *β* regarding the fixed value of *α*). The 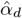 and 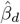 estimators are used for simulations, instead of 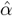 and 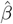 estimators, because number of cells, mass and volume of solid tumors are measured discretely over time in *in vitro* [24], preclinical [1–5, 7, 15] and clinical [25] studies.

The results of this study reveal the existence of *β*_*lim*_ and *σ*_*lim*_, which may suggest that the unperturbed solid tumor is active (exists and functional: *β* ≤ *β*_*lim*_ and *σ* ≤ *σ*_*lim*_), inactive (exist and non-functional: *β* ≤ *β*_*lim*_ and *σ > σ*_*lim*_, or *β > β*_*lim*_ and *σ* ≤ *σ*_*lim*_) or does not exit (*β > β*_*lim*_ and *σ > σ*_*lim*_), as is explicitly evidenced in Figures 5 and 6 (unprecedented in the literature). Futhermore, this study shows how the degree of stochasticity inherent in unperturbed TGK is unfluenced by the parameters *V*_*obs*_, *σ*, and the difference of *α* respect to *β* (*α > β*), keeping constants *α* and *V*_0_ may be estimated from experimental data following the methodology reported in [4, 6, 7].

Ferrante et al. [15] report that 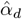 tends to *α* when *T* increases; nevertheless, T is limited in preclinical studies by bioethical guidelines (tumor burden should not usually exceed 10 % of the host animal’s normal body mass) [6, 7]. This why, the results in Figures 1-4 are only shown for tumor volume ≤ 5 cm3 and time ≤ 30 *days*, as observe in our previous experimental studies [3, 4, 6, 7]. Consequently, insufficient data resources in cancer may appear in the literature. Therefore, the practical utility of existing parametric approaches for identifying deterministic and stochastic differential equations, as both versions of MBBG equations, is frequently limited.

The analysis of stochastic MBBG equation is a challenge because it allows us to understand the inherent stochasticity of data and the complexity of dynamical systems, as the unperturbed cancer [26]. This stochasticity occurs from early stage of tumorigenesis by intrinsic (in whole cancer) and external (in cancer environment or surrounding healthy tissue) endogenous/physiological noises generated by different sources. These noise sources are interconnected, happen on different space-time scales, and influence markedly on cancer dynamics and complexity, and participate in cancer in cancer cells proliferation, progression, metastasis, anti-cancer therapy resistance, and non-recognition of the immune system [8, 27, 28].

Among intrinsic endogenous noise sources in unperturbed solid tumors may be mentioned: 1) cell-cell and cell-extracellular matrix interactions involved during entire unperturbed TGK (complex and dynamic process) [8]; 2) the 14 clinical hallmarks of cancer themselves [29]; 3) intratumor heterogeneity [30–32]; stress due to differences in metabolic demands among cancer cells [30–32]; 4) point mutations present stochastically in high heterogeneous tumors [32, 33]; 5) dynamic formation of vacancies (e.g., oxygen vacancies) in cancer that cause instabilities in space-time during tumor growth [3]; 6) dynamical structural transformation that happens in the entire TGK [3], among other possible unknown noise sources. Nevertheless, some external endogenous noise sources in unperturbed solid tumors may influence in TGK, such as: 1) interactions among noncancerous and cancerous cells in the tumor environment [32]; 2) environmental noise [10, 34]; 3) body temperature [3, 35], among others.

The hallmarks of cancer are sustaining proliferative signaling, evading growth suppressors, inducing or accessing vasculature, deregulating cellular metabolism, avoiding immune destruction, non-mutational epigenetic reprogramming, polymorphic microbiomes, senescent cells, unlocking phenotypic plasticity, enabling replicative immortality, resisting cell death, activating invasion and metastasis, genome instability and mutation, and tumor-promoting inflammation [29]. These hallmarks are involved in intratumor heterogeneity (cell clones that differ in their genotype and/or phenotype, and mechanical and electrical properties) [17, 30–32]. Intratumor heterogeneity is a consequence of the combination of extrinsic factors (from the tumor environment) and intrinsic parameters from the cancer cells themselves (e.g., genetic, epigenetic and transcriptomic traits; ability of proliferation, migration and invasion; stemness and plasticity) [3, 31, 36].

The itself mutations that take into account in cancer during its growth are considered as random process [33] and contribute to deregulated signaling mediators that lead to increased genomic instability, tumorigenic transformation, among others alterations [37]. Interactions among noncancerous and cancerous cells in the tumor environment may contribute to the diversification of cancer forces [32]. These complex interactions may be mechanical [3–5, 18], electrical [17], chemical [16, 38] and biological [32] in nature. Furthermore, the environmental noise to the tumor (coming from the surrounding healthy tissue and/or from outside the human body) may influence in stochastic tumor-immune system [10, 34], which also leads to tumor stochastic behavior.

The different noise types are considered in several mathematical models, such as: prey-predator-based model [8], both white noise and Markov switching [38], stochastic Lyapunov analysis and formula of 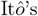 [34], multiplicative non-Gaussian colored noise [40], and non-linear mixed effects [8] and noise-induced dynamic (unavoidable random fluctuacions) in the cancer [27].

Although entire TGK and biophysical-chemical-metabolic processes in cancer occur at constant temperature in the organism, the thermal energy associated with this temperature constitutes a noise source for the cancer formation and growth in this host [3]. Lim et al. [35] hypothesize the existence of an external noise from thermal fluctuations of local ligand concentrations to understand chemotaxis of cancer cells. These thermal fluctuations originate random errors in binding reactions and signal transduction reactions due to the asymmetrical cellular distribution of substrate-bound receptors to recognize a chemical gradient. For this, they use a mathematical model based on signal/noise ratio that represents cancer stochastic properties.

The stochasticity in untreated solid tumors above-mentioned may explain why they exhibit an oscillatory behavior with non-constant frequency probably by generation of imbalance states, as report in different studies [8, 36]. These imbalance states may justify dynamical structural transformations that happen in the entire unperturbed TGK [3]. The oscillatory behavior in cancer may be related to the stochastic resonance phenomenon, aspect verified with a stochastic tumor growth model under the excitation of Lévy noise and Gaussian white noise, which are useful to simulate environmental disturbances [28]. Hu et al. [14] demonstrate that Lévy jumps noise, the white noise and switching noise influence importantly on the properties of the hybrid stochastic delay Gompertz model.

As fluctuating environment alters enzymatic reactions of proteins in solid cancer, as a biochemical system, as solid tumors, we incorporate white noise in the stochastic MBBG equation. Das et al. [41] report that this environmental noise may have an important role in processes of stochastic permanence and extinction of tumor cells. Nevertheless, intrinsic and external noises cannot be analyzed separately, otherwise they overlap. Consequently, the cancer senses a net noise, which multiplies over time during tumor growth. These mixed noises may be included in stochastic differential equations that allow describing stochastic hybrid [40], mixed processes that happen stochastically [42], and population mixed-effects with stochastic version of Norton-Simon-Massague tumor growth model [43], but using the stochastic MBBG equation for tumor growth, as report in this study.

The results in Figures 2-6 and existence of *β*_*lim*_ and *σ*_*lim*_ confirm some findings reported in the literature, such as: the existence of limit parameters for stochastic permanence of cancer cells [41]; the Allee effect happens in the entire unperturbed TGK [9, 45–47]; Wiener process (or Brownian motion) [44], and three possible functional states of the solid tumor (active, inactive or non-exist) [4].

The stochastic permanence of cancer cells is related to the oscillation of them [41], aspect that may be verified in this study by the high dispersion observed in the entire TGK with the increase of *V*_*obs*_ and *σ*, and decrease of the difference of *β* respect to *α*, provided that *β* ≤ *β*_*lim*_ and *σ* ≤ *σ*_*lim*_.

The Allee effect is a weak or strong positive relationship between the overall individual fitness (e.g., cancer cells) and population size/density (e.g., solid tumor). Its apparition mechanism is inherently related to survival and reproduction from aggregation, cooperation or facilitation among cancer cells in the solid tumor that permit them to growth, invade and metastasize more efficiently, and defense against immune system attack and anticancer therapies [9, 45–47].

The weak Allee effect (reduced positive *per capita* growth rate directly link to individual fitness of the cancer) is exhibited at lower solid tumor density/size, which the cooperation is generally ineffective [48]. This may be explained for two possible reasons. First, nucleation and aggregation mechanisms happen slowly during the avascular phase of unperturbed TGK (first stage of this kinetics, which is comprised from the moment of inoculation/induction of cancer cells in the organism until the moment in which the tumor reaches a certain volume from which unperturbed TGK is triggered, named *V*_*triggered*_). Second, mutuations and duplication of cancer cells are unfavorable in this avascular phase of TGK and characterized by instabilities due to fluctuations and high diffusion [49], in agreement with Johnson et al. [47], who explain both Allee effect and stochasticity in the cancer cell growth at low density from diffusion process in solid tumors. Our previous experience in mice bearing of solid tumors suggest that these two processes occur for tumor volumes *<* 0.1 *cm*^3^ depending on the histological variety [3, 4, 6, 7, 49].

The fluctuations and high diffusion at avascular phase of the unperturbed TGK agree with the statement that the elimination of the cancer has a higher probability at this phase that vascular phase at unperturbed TGK (larger solid tumors: *V* ≥ *V*_*triggered*_), as report in previous studies [9, 45–48]. In order to avoid self-destruction of the solid tumor at avascular phase of unperturbed TGK due to environmental fluctuations, and those inherent in the solid tumor itself, the cancer emerges the vascular phase of TGK [3] to guarantee high growth rate (increase of tumor size) for the cooperation among cancer cells [9, 45–48] and production of pro-angiogenic factors (e.g., vascular endothelial growth factor) that creates blood vessels to irrigate the cancer for its growth, invasion and metastasis [50].

Different researchers report that the solid tumor exhibits a strong Allee effect at vascular phase from a critical density/size under which the cancer growth rate becomes negative. In this case, the solid tumor tends to disappear without any further aid because the cancer density/size hits a number below this threshold [9, 45–47]. This agrees with the results of this study for *β > β*_*lim*_ and *σ > σ*_*lim*_ (see black square for each curve in Figures 5 and 6).

The condition *β > β*_*lim*_ may mean an overproduction of endogenous antiangiogenic molecules (e.g., endostatin, angiostatin, thrombospondin) that breaks and dominates the balance with the production of proangiogennic molecules (favor the angiogenesis process) and/or prevails over cancer cells duplication (*η* ≥ *α*, keeping constant the *α* value). In both cases, the cancer self-destruction and disappears physiologically without any further aid, in agreement with Castañeda et al. [4]. In the first case, neovascularization and metastasis processes are physiologically inhibited. If neovascularization is inhibited, the solid tumor not receive nutrients and oxygen; therefore, it dies. In the second case, the overproduction of endogenous antiangiogenic molecules not only dominates the duplication process of cancer cells, but may activate cell loss mechanisms (necrosis, apoptosis, metastasis and exfoliation) [51] to compensate this exaggerated antiangiogenic process.

The condition *σ > σ*_*lim*_ may suggest exaggerated diffusion of several components in the unperturbed cancer (e.g., cancer cells, molecules, ions, electrons) that lead to very fast movements of them in the cancer, which becomes highly unstable and self-destructs. This may be explained from biophysical point of view because the cancer does not have favorable conditions for their growth, invasion and metastasis. In this case, marked instabilities in cancer may induce also high instabilities in the surrounding healthy tissue and the rest of the organism that may lead to irreversible changes in it or to its death. The instabilities on both tissues may induce high fluctuations with respect to the physiological fluctuations that are permissible for the organism.

We do not rule out the condition *β > β*_*lim*_ and *σ > σ*_*lim*_ correspond to the marked depletion and cachexia of the organism with cancer observed in mice (tumor volume and mice weight decrease from tumor volumes larger than 6 *cm*^3^) [52–54] and humans (Karnofsky performance status (≤ 30 %) [55] or ECOG performance status (≥ 4) [56]). Nevertheless, ethical aspects in tumor-bearing animals (tumor volume does not exceed 10 % of the weight) do not allow observing these two conditions [3, 4, 6, 7].

The inequalities established for *β*_*lim*_ and *σ*_*lim*_ should not be analyzed separately because they are interconnected with each other and with the parameter *α*. This interconnection defines three possibles states for unperturbed solid tumors depending on the values of *β* and *σ*: active (exists and functional: *β* ≤ *β*_*lim*_ and *σ* ≤ *σ*_*lim*_), inactive (exists and non-functional: *β* ≤ *β*_*lim*_ and *σ > σ*_*lim*_, or *β > β*_*lim*_ and *σ* ≤ *σ*_*lim*_) or does not exist (*β > β*_*lim*_ and *σ > σ*_*lim*_), as is explicitly evidenced in Figures 5 and 6. These three unperturbed tumor states ares also documented when the analysis of TGK is made form the fractal dimension of the tumor countour (*d*_*f*_) [4]. Castañeda et al. [4], report that the cancer may only exist for *α* values that correspond to 0 *< d*_*f*_ *<* 1; nevertheless, the cancer is non-functional or self-destructs for 1 *< d*_*f*_ *<* 1.5 and *d*_*f*_ *>* 1.5, respectively. The connection of these three states of this study and those in [4] may be explained because *β* and *σ* depend on *α*, which in turn depends on *d*_*f*_ ; therefore, *β* and *σ* depend also on *d*_*f*_.

The active tumor state (1 *< d*_*f*_ *<* 1.5, *β* ≤ *β*_*lim*_ and *σ* ≤ *σ*_*lim*_) confirm different findings reported in the literature, such as: 1) the sigmoidal growth that exhibit unperturbed solid tumors when they are induced in the organism [1–4, 6, 7]; 2) the angiogenesis is an emergent, regulated and self-organized process [3, 4]; 3) the cooperativity, self-organization, self-similarity and coherence in cancer during entire unperturbed TGK [3, 4, 17, 48, 49, 57, 58]; 4) the vascular network in cancer forms patterns in space-time and the fractal dimension of the vascular branching architecture over time displays a sigmoidal pattern [59]; 5) the balanced and regulated production between proangiogenic and antiangiogenic molecules during entire unperturbed TGK [60]; 6) the solid tumor is coherent, self-organized, self-similar and efficient [3, 4, 6, 57–59]; 7) the cancer is constantly renewed (cancer passes to a new state during its entire TGK), in agreement with other studies [3, 49, 61]; and 8) the physiological anti-angiogenesis and diffusion mechanisms are limited and self-regulated during the growth of unperturbed solid tumors [3, 4].

The limit conditions for *β* and *σ* suggested in this study (*β* ≤ *β*_*lim*_ and *σ* ≤ *σ*_*lim*_) and 1 *< d*_*f*_ *<* 1.5 [4], as well as the sigmoidal behavior of tumor fractal dimension [59], and Allee effect mechanisms [9, 45–47] may have an essential role in cancer growth and in to explain why the entire unperturbed TGK does not follow an exponential growth, but sigmoidal, as widely documented in experiment [1–4, 6, 7, 15, 52–54, 59]

The point of coordinates (*β*_*T*_, *σ*_*T*_) in the unperturbed TGK (see black squares in 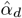 versus 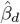 graph, Figure 6) delimits its growing part (*β* ≤ *β*_*T*_ and *σ* ≤ *σ*_*T*_, corresponding to an active solid tumor) from decreasing part (*β > β*_*T*_ and *σ > σ*_*T*_, corresponding to an inactive solid tumor). A peak is observed on the growing part of TGK (*β* ≤ *β*_*T*_ and *σ* ≤ *σ*_*T*_) whose morphology (width and amplitude) and positioning depend on *β, σ* and *V*_*obs*_ for *α* constant. This peak is characterized because 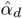 first decreases up to reach its minimum value and then increases when 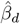 increases up to the point (*β*_*T*_, *σ*_*T*_). This peak may be related to the transition between avascular and vascular phases of unperturbed TGK and/or stochastic resonance in this kinetics.

The transition between avascular and vascular phases of unperturbed TGK is observed in preclinical studies for *V*_*obs*_ *<* 0.5 *cm*^3^ [3, 4, 6, 7, 49]. Nevertheless, the results in 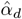 versus 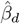 graph may suggest that this transition may occur for any tumor volume greater than Vobs and less than 0.5 *cm*^3^, most likely when the solid tumor volume is *V*_*triggered*_, in agreement with the peak that is observed in plot of local Avrami exponent versus *ln*(*t*) [3]. This may be explained because the coefficients of MBBG and KJMA equations have been analytically related to *α* and *β* [5].

The sharper peak (narrower width and higher amplitude) that is observed in 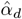 versus 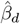 graph for the lowest values of *σ* and *V*_*obs*_ may correspond to smallest solid tumors (tumor volumes much smaller than *V*_*triggered*_) with the ability to rapidly duplicate and metastasize their cells. Therefore, they reach the point (*β*_*T*_, *σ*_*T*_) in the shortest time. To avoid TGK from being non-exponential, the unperturbed tumor must produce more endogenous antiangiogenic molecules, aspect that may explain why the position of the peak is further to the right for tumors with the lowest values of *σ* and *V*_*obs*_, in agreement with Castañeda et al. [4]. In this case, the avascular phase of the unperturbed TGK of these solid tumors first happens slowly (heterogeneous nucleation mechanisms governed by diffusion are slower) and then very quickly in the time (growth and interaction mechanisms governed by diffusion pare faster), in agreement with González et al. [3] and Cabrales et al. [49].

The mentioned in the previous paragraph is not observed when *σ* increases for the lowest value of *V*_*obs*_ because the peak is wider, lower amplitude and runs further to the left; therefore, the tumor volume reaches the point (*β*_*T*_, *σ*_*T*_) in the longest time. This finding is evident when *V*_*obs*_ increases, being marked for *V*_*obs*_ = *V*_0_. This situation may correspond to solid tumors mensurable in the experiment, whose tumor volumes are close (below) or greater (but ≤ *V*_0_) than *V*_*triggered*_. In this case, the avascular phase of the unperturbed TGK of these solid tumors first happens rapidly (heterogeneous nucleation mechanisms governed by diffusion are faster) and then very slowly in the time (growth and interaction mechanisms governed by diffusion are slower), in agreement with González et al. [3] and Cabrales et al. [49].

The discussed in two previous paragraphs may suggest that solid tumors that have rapid at avascular phases are those that grow more slowly at the vascular phase and vice versa. This hypothesis can be demonstrated directly from the results of González et al. [3]. An additional theoretical study will be carried out to demonstrate this hypothesis.

In 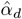 versus 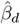 graph, the broadening of the peak when *σ* and *V*_*obs*_ increase may be explained from different findings reported in the unperturbed cancer, such as: greater production and mobility of cancer cells [20, 62]; charged carriers (e.g., positively and negatively charged ions, electrons) [17, 63]; different types of molecules (e.g., angiogenic molecules, reactive oxygen and nitrogen species, and free radicals) [64, 65]; and distinct family of high mobility group proteins (e.g., HMGI-C, HMGI(Y)) [66]. These elements may participate in the complex network of signaling and interactions; greater heterogeneity, anisotropy, compactness, elasticity, fluidity, amorphous characteristics, deformation and irregular borders of the solid tumors [3, 4, 6, 7, 20, 57–59, 67, 68], as well as in their altered bioelectricity and dysregulated metabolism [17, 63].

On the other hand, resulting peak broadening may be consequence of the combination of all individual broadenings during unperturbed TGK due to the high heterogeneity and anisotropy in solid tumors. This may be explained because the solid tumor is composed of cellular clusters with different genotypic and phenotypic characteristics, and electrical properties [3, 4, 17, 49]. Furthermore, broader peak may be a measure of defects in the tumor mass, dynamic transformations, anisotropic broadening and pressure broadening in the unperturbed TGK. All these aspects have an essential role in the growth, progression and metastasis of the unperturbed cancer [17, 63, 69]. The anisotropic broadening is corroborated in experimental [3, 4, 6, 7] and theoretical [20] studies. The pressure broadening may be relate to interstitial fluid pressure in solid tumor growth that lead to heterogeneous mechanical stress [70, 71].

The increase of the peak broadening and its change of position in 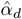 versus 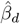 graph when *V*_*obs*_ increases may be explained by the compactness of the solid malignant tumors due to the packaging of their cells (cancer cells very close together) into a solid mass [3, 4, 6, 7]. This may corroborate the weak interactions among them during solid tumor growth, being marked for the most values of *V*_*obs*_ because more cells are produced, as report in [3, 4, 20]. Previous studies suggest that the nature of these interactions is electrical, which has been associated with the [17, 72, 73]. Furthermore, the broad peak is this peak broadening may be an indicator of irreversible biophysical-chemical processes in the tumor, being marked for the highest value of *V*_*obs*_. This aspect may be related to the low effectiveness of any anticancer therapy for large tumor sizes, both in preclinical and clinical studies.

The discussed above may suggest why broadened peaks do not take the form of a Gaussian distribution (inhomogeneous broadening of each peak), which corroborates the inherent fluctuactions coming from various noise sources in different parts of the tumor and its environment during the entire TGK when *σ* and *V*_*obs*_ increase, being marked for the higher values *V*_*obs*_. This broaden unevenly may lead to an irregular and asymmetric shape of each peak.

The inhomogeneous broadening of each peak may be an indicator of a stochastic transition on the unperturbed TGK probably caused by various cell clusters interacting with each other and respond in a different way to external perturbations (or have varied interactions with their surroundings). We hypothesize that this may be possible because these clusters have dissimilar different frequencies and transition energies since they have different genotypic and phenotypic characterisitcs, and electrical properties. This confirms that inhomogeneous broadening is frequently observed in complex systems, as the unperturbed solid tumors, in which heterogeneity of energy levels and local dynamical changes (due to autoorganizations by point defects in the solid tumor mass) happen, corroborating that unperturbed TGK is a dynamical transformations kinetics [3]. Furthermore, we do not discard that electric fields and electric field gradients from charged defects and/or carriers (e.g., electrons, ions) [17, 63] may lead to this inhomogeneous broadening of peak.

## New insights

Knowledge of the inequalities established for *β* and *σ* (results of this study) and *d*_*f*_ [4] stimulates the establishment of analytical or numerical relationships to know the parameter space (space of possible parameter values of *β, σ* and *d*_*f*_) for which the unperturbed solid tumor is functional (forms, grows and invade nearby and distant tissues), non-functional or self-destructs.

A further study can be carried out to know how other noise types (e.g., Lévy noise [14, 28] or another type of non-Gaussian noise [74]), different to Gaussian white noise, affect the dynamics of the unperturbed TGK described with the stochastic MBBG equation (Equation (5)) for the likelihood estimators of *α* and *β* for both continuous (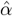 and 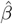) and discrete (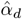 and 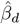) cases shown in this study. This is taken into account that Gaussian white noise and Lévy noise have different effects on the cancer dynamical behavior [28].

A fractional stochastic MBBG equation may be proposed to describe simultaneously high heterogeneities and multiscale fluctuations (due to intrinsic and external noises) that exhibit unperturbed solid tumors. This is taken into account that systems with these two characteristics can be described with fractional differential equations [75, 76].

The ideas of this study may be extended to that reported in [3] to know the stochastic version of the modified Kolmogorov-Johnson-Mehl-Avrami (mKJMA) equation and its respective likelihood estimators *K, n* and *λ* for both continuous and discrete cases. These likelihood estimators can be related analytically or numerically to those of *α* and *β* reported in this study for both continuous (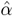 and 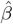) and discrete (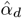 and 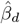). For this, a similar methodology to that documented in [5] must be followed.

Factors that influence in the cancer degradation processes should be included in Equation 5, resulting a new Wiener-process-based model, to achieve an accurate estimation of experimental data in cancer. For this, the study of Zhang et al. [61] should be taken into account. This is important because environmental fluctuations are omnipresent in the cancer cells. Therefore, the cancer does not follow deterministic laws strictly, as report in [1–6, 20]. Also, the deterministic model does not provide any insight into the likelihood of extinction of the cancer and other disease types. These reasons justify why we use the stochastic MBBG in this study.

An external perturbation (e.g., electrochemical therapy or electrolytic ablation, named EChT [6, 7]) may be added in the stochastic MBBG equation (Equation 5), as in [20], to know how the explicit expressions of likelihood estimators of *α* and *β* for both continuous (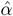 and 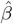) and discrete (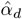 and 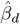) cases for EChT perturbed tumors and how these estimators depend on cancer biological characteristics and parameters of this physical therapy. In principle, this idea may be extended to any anti-cancer therapy.

## Conclusion

The stochastic Montijano-Bergues-Bory-Gompertz equation may be applied in the experiment to describe the unperturbed tumor growth kinetics, as previously demonstrated for its deterministic version, in order to estimate the parameters of this equation and their connection with processes involved in the growth, progression and metastasis of unperturbed solid tumors.

